# PRDM paralogs are required for Meckel’s cartilage formation during mandibular bone development

**DOI:** 10.1101/2025.07.17.665392

**Authors:** Qootsvenma Denipah-Cook, Bryanna V. Saxton, Kristin B. Artinger, Lomeli C. Shull

## Abstract

Mandibular bone development utilizes both endochondral ossification, forming from the cartilaginous anlage Meckel’s cartilage derived from neural crest cells (NCC) and intramembranous ossification with direct differentiation of NCCs toward osteoblasts. Wnt/β-catenin signaling drives osteogenic vs chondrogenic differentiation and must be tightly controlled during the differentiation of osteochondroprogenitors. Chromatin remodelers add hierarchal regulation to the activation and repression of crucially timed gene regulatory networks and signaling cascades. In this study, we investigated the function of two chromatin remodelers—histone methyltransferases, PRDM3 and PRDM16 during murine craniofacial development. Conditionally ablating both *Prdm3* and *Prdm16* in the neural crest lineage using the Wnt1-Cre driver resulted in dramatic craniofacial phenotypes, including a severely hypoplastic mandible with complete absence of Meckel’s cartilage at E18.5. Focusing on the Meckel’s cartilage and mandibular bone phenotype, histological analysis demonstrated a significant increase in RUNX2+ osteoblast precursors, and loss of SOX9+ chondrogenic cells, suggesting an increase in osteoblast progenitors at the expense of chondrocytes that would otherwise form the Meckel’s cartilage. This was not due to alterations in proliferation or apoptosis, as we observed no significant changes in the number of phosphoH3+ or cleaved caspase3+ cells in the mandibular process at E11.5, suggesting lack of NCC-derived chondrocytes is due to a change in NCC osteochondroprogenitor fate decisions. mRNA transcripts and protein abundance of Wnt/β-catenin signaling components were elevated in the mandibular process during initial NCC osteochondroprogenitor condensation events, suggesting PRDM3 and PRDM16 normally restrict expression of Wnt/β-catenin signaling components during NCC-derived osteochondroprogenitor differentiation to promote chondrogenesis and Meckel’s cartilage formation. Taken together, PRDM3 and PRDM16 are required for NCC differentiation toward chondrocytes during Meckel’s cartilage formation by controlling proper spatiotemporal Wnt/β-catenin transcriptional activity and this process is necessary for morphogenesis of the developing mandible.

## Introduction

Neural crest cells (NCCs) are a multipotent population of cells that have the unique ability to give rise to a variety of cell types and tissues throughout the body. NCCs are specified at the neural plate border during neurulation. As the neural tube begins to fold, NCCs will undergo an epithelial to mesenchymal transition, delaminate from their extracellular matrix and become a highly migratory population of cells^1–5^. NCCs travel long distances along the anterior-posterior axis of the developing embryo and when they reach their destination, begin to differentiate toward a variety of specific cell types. The NCCs that originate in the midbrain and hindbrain rhombomeres from the lateral ridges of the neural plate will migrate in streams to populate the pharyngeal arches and frontonasal mass^1,2,6^. These NCCs, termed cranial neural crest cells, will differentiate into a variety of cells that contribute to the formation of the craniofacial complex, including chondrocytes, osteoblasts, odontoblasts of the teeth as well as neurons and glia of the peripheral nervous system. There are numerous transcription factors and gene regulatory networks (GRNs) that facilitate the establishment of specific cranial neural crest cell fate lineages, however the molecular mechanisms underlying how cell fate decisions are spatiotemporally controlled during the formation of the craniofacial complex remain incomplete.

There are two types of bone formation that contribute to the formation of the craniofacial complex: 1) intramembranous ossification, or direct differentiation of a progenitor mesenchyme toward the osteoblast lineage, and 2) endochondral ossification, where condensed mesenchyme differentiates toward the chondrocyte lineage to form a cartilaginous anlage that serves as a ‘template’ for subsequent bone formation. Most bone structures that comprise the craniofacial skeleton are dermal and formed through intramembranous ossification. There are some cranial bones that use endochondral ossification, including the structures of the cranial base (presphenoid, basisphenoid and basioccipital). The mandible, however, utilizes both intramembranous and endochondral bone formation, whereby condensed neural crest mesenchyme in the first pharyngeal arch give rise to chondrocytes forming long bi-lateral struts of cells on either side of the mandibular process that meet and fuse anteriorly^7^. This transient cartilaginous structure, termed Meckel’s cartilage, is crucial in the morphological development and outgrowth of the mandible. Meckel’s cartilage also contributes to the formation of the anterior mandibular symphysis, the posterior middle ear ossicles (malleus and incus) and the anterior ligament of malleus and sphenomandibular ligament^8^. While the diverse fates of Meckel’s cartilage and the GRNs and signaling pathways that drive chondrogenesis vs osteoblastogenesis are more defined, less is known about the factors controlling the initial steps of cranial neural crest commitment toward chondrocytes in the mandibular process in order to facilitate the formation of Meckel’s cartilage.

Several epigenetic regulators and chromatin remodelers have important roles in spatiotemporally controlling chromatin accessibility and subsequently gene expression of GRNs during cranial neural crest development and formation of craniofacial derivatives and structures^9^. Among these are the PRDM family of lysine methyltransferases. This family of methyltransferases has the unique ability to control gene expression through several mechanisms including epigenetically modulating chromatin accessibility, directly binding DNA via zinc-finger domains, or interacting with other protein complexes^10–12^. While there are 17 different members of the PRDM family in the human genome, genome wide association studies have associated single nucleotide polymorphisms in the genes encoding two of these PRDMs, PRDM3 (EVI1/MECOM) and PRDM16, with craniofacial abnormalities, including cleft lip/palate and variation in facial morphology^13–19^. Numerous studies have implicated the importance of PRDM3 and PRDM16 in key developmental processes, including craniofacial development, neurogenesis, and lung morphogenesis^13,20–22^. In zebrafish, Prdm3 and Prdm16 control craniofacial chondrocyte differentiation by antagonistically balancing Wnt/β-catenin transcriptional activity through modulating chromatin accessibility of putative Wnt/β-catenin target genes. Recently, others have also shown that PRDM3 and PRDM16 together mediate murine alveolar epithelial cell differentiation during lung morphogenesis by altering chromatin accessibility around target NKX2-1 genes^22^. Mouse models have shown neural crest specific ablation of either *Prdm3* or *Prdm16* causes craniofacial defects^13,20^. Homozygous *Prdm3* mutant animals exhibit more subtle phenotypes, namely mild anterior mandibular hypoplasia and marginal defects in the extension of their snout^20^. *Prdm16* homozygous mutant animals present with a range of craniofacial abnormalities including snout extension defects, anterior mandibular hypoplasia, clefting of the secondary palate and severe hypoplasia of the middle ear ossicles (tympanic ring, incus, malleus, and retroarticular process of the squamosal bone)^20^. The shared phenotype of mandibular hypoplasia in both *Prdm3* and *Prdm16* single mutant animals is associated with altered chondrocyte organization in the developing Meckel’s cartilage of these animals, suggesting each factor has a particular functional role in coordinating chondrogenesis. The combined function of both PRDM3 and PRDM16 in mediating the development of Meckel’s cartilage chondrocytes remains unknown.

In this study, we investigated the genetic interaction between *Prdm3* and *Prdm16* during mammalian neural crest development and formation of the craniofacial structures by ablating both *Prdm3* and *Prdm16* in the neural crest lineage using the Wnt1-Cre driver. Loss of both *Prdm3* and *Prdm16* leads to a more hypoplastic lower jaw due to complete loss of the Meckel’s cartilage. In their absence, expression of Wnt/β-catenin signaling components remain elevated and promotes the differentiation of NCC-derived osteochondroprogenitors toward the osteoblast lineage at the expense of NCC-derived chondrocytes that would otherwise form Meckel’s cartilage. We hypothesize that PRDM3 and PRDM16 control cranial neural crest cell fate decisions in the mandibular process by restricting expression of Wnt/β-catenin signaling components during NCC-derived osteochondroprogenitor differentiation programs to allow for chondrogenesis and formation of Meckel’s cartilage. Through controlling Meckel’s cartilage development, these chromatin remodelers ultimately contribute to proper mandible morphology and formation of the middle ear ossicles.

## Results

### Combined loss of *Prdm3* and *Prdm16* in the neural crest lineage causes craniofacial defects

To assess the genetic interaction between *Prdm3* and *Prdm16* during murine neural crest development, we conditionally ablated both *Prdm3* and *Prdm16* in the murine neural crest lineage using the Wnt1-Cre driver^23^. Homozygous *Prdm3^fl/fl^;Prdm16^fl/fl^;Wnt1-Cre*^+/Tg^ mutant animals were present at Mendelian ratios and survived to embryonic day (E) 18.5, but exhibited more severe phenotypes compared to each single mutant alone. Viability of double mutant embryos was not carried out past E18.5. However, we predict that they would not survive long after birth given the severity of their phenotypes present at E18.5, the phenotypes of single mutant animals^20^, and the functions of these factors in heart (*Prdm3*) and cleft palate development (*Prdm16*)^13,15,16,24,25^. Gross anatomical assessment of *Prdm3^fl/fl^;Prdm16^fl/fl^;Wnt1-Cre^+/Tg^* double mutants at E18.5 revealed a variety of craniofacial defects, including abnormal snout extension, very low set pinnae and mandibular hypoplasia (**Fig 1A-1B**). MicroCT 3D reconstructions further exemplified the craniofacial defects observed in double mutants, with severe shortening of the maxilla, fusion of the premaxilla and maxillary bones, complete loss of middle ear ossicles, and loss of the retroarticular process of the squamosal bone and occipital bone (**Fig 1D-1E**). These morphological defects were also assessed by Alcian blue and Alizarin red stained whole mount skeletal preparations (**Fig 1H-1I**) and were more drastic than single mutants (**Supplemental Fig 1A-C’**).

**Figure 1.**
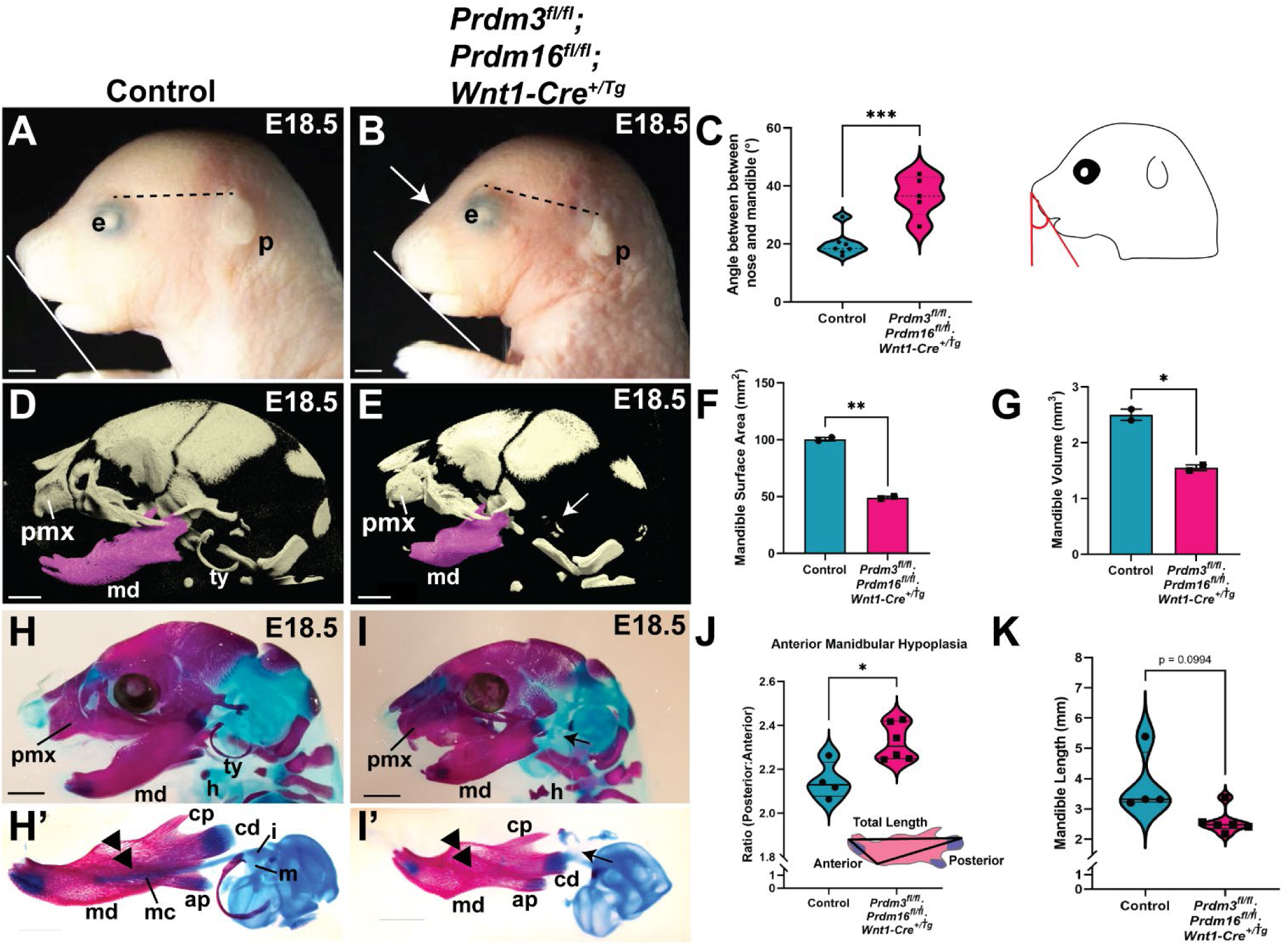
Ablation of both *Prdm3* and *Prdm16* in the neural crest lineage causes craniofacial defects. (A-B) Lateral images of the head show snout defects (white arrows), mandible hypoplasia (white lines), and low set pinnae in *Prdm3;Prdm16* double mutants (indicated by dotted black line drawn from the top of the eye to the top of the pinna). (C) Quantification of mandibular hypoplasia was performed by measuring the angle generated with a line drawn from the snout to the mandible (n=7 for control; n=5 for double mutant). (D-E) 3D reconstructed lateral images from microCT scans of the head show cranial defects in double mutants compared to controls. Segmented mandible in purple. (F, G) Surface area (F) and volume of the segmented mandibles (G) from the microCT reconstructions (n=2 for control and double mutant). (H-K) Alcian Blue and Alizarin Red stained whole mount skeletal preparations at E18.5. (H’-I’) Alcian blue and Alizarin red stained mandibles were dissected and removed from control or *Prdm3;Prdm16* double mutants. Shown are lateral views of the mandible. Black double arrow heads indicate where Meckel’s cartilage is present in control animals but completely absent in *Prdm3;Prdm16* double mutants. (J-K) Quantification of anterior mandibular hypoplasia was performed by measuring the ratio of the length of the anterior portion of the mandible relative to the posterior length (J) and the overall mandible length (mm) (K) (n=4 for control; n=6 for double mutant). Abbreviations: ap, angular process; cd, condylar process; cp, coronoid process; h, hyoid; i, incus; m, malleus; mc, Meckel’s cartilage; md, mandible; pmx, premaxilla; ty, tympanic ring. \**P≤*0.05, \*\**P≤*0.005, \*\*\**P≤*0.0005 (unpaired, two-tailed Student’s *t* test).

We previously showed that *Prdm3* and *Prdm16* single mutants each develop mild anterior mandibular hypoplasia, a likely consequence of abnormal chondrocyte differentiation and maturation in the Meckel’s cartilage^20^. Given this shared phenotype across both single mutants and the increased severity of this phenotype in the double mutants, we focused on mandibular development in the double mutant animals (**Fig 1H’-I’**). The mandible in *Prdm3^fl/fl^;Prdm16^fl/fl^;Wnt1-Cre^+/Tg^*double mutants was notably smaller and morphologically different compared to littermate control animals at E18.5. This mandibular hypoplasia was quantified by assessing the angle between the end of the snout and the mandible, which was significantly larger in double mutants compared to controls (**Fig 1C**). Additionally, the ratio between the anterior portion of the mandible relative to the posterior length was significantly greater in *Prdm3^fl/fl^;Prdm16^fl/fl^;Wnt1-Cre^+/Tg^*double mutants (**Fig 1J**). Overall mandibular length was dramatically smaller (**Fig 1K**), as was the mandible surface area (**Fig 1F**) and volume (**Fig 1G**) relative to controls as quantified from dissected Alcian blue and Alizarin red stained whole mount skeletal preparations and MicroCT 3D reconstructions, respectively. In E18.5 control animals, the Meckel’s cartilage can be seen extending from the middle ear ossicles toward the middle domain of the mandible (**Fig 1H’**) in Alcian blue and Alizarin red whole mount skeletal stains. Notably, in *Prdm3^fl/fl^;Prdm16^fl/fl^;Wnt1-Cre^+/Tg^*double mutants, the Meckel’s cartilage is completely absent (**Fig 1I’**) and cartilaginous middle ear ossicles are either missing or severely hypoplastic (**Fig 1E, I’**). This striking phenotype is highly specific to only the Meckel’s cartilage as other endochondral bones form, such as the cranial base (presphenoid, basisphenoid and basioccipital) and nasal cartilages (**Supplemental Fig 2**). Absence of Meckel’s cartilage and middle ear ossicles in double mutants suggests endochondral ossification bone formation during mandible development is compromised.

To test whether intramembranous ossification was affected in the developing mandible, von Kossa staining was performed on coronal sections through the mandible at E15.5 to assess mineral deposition (**Supplemental Fig 3**). There were no obvious defects in mineralization observed in the anterior portion of the mandible of double mutants (**Supplemental Fig 3A-B**). More posteriorly, the area of deposited bone appeared marginally thicker in double mutant animals compared to controls, and as observed at E18.5, Meckel’s cartilage was completely absent throughout the anterior-posterior axis of the mandible in *Prdm3^fl/fl^;Prdm16^fl/fl^;Wnt1-Cre^+/Tg^*mutants (**Supplemental Fig 3A’-B’**). Importantly, the trabecular networks that form through intramembranous ossification are comparable to those observed in control animals, indicating that intramembranous bone formation was not affected in *Prdm3^fl/fl^;Prdm16^fl/fl^;Wnt1-Cre^+/Tg^*double mutants. To test if the lack of Meckel’s cartilage at E15.5 was the result of increased infiltration of osteo/chondroclasts responsible for resorption of the intermediate portion of Meckel’s cartilage anlage, tartrate resistant acid phosphatase (TRAP) staining was performed (**Supplemental Fig 3**). Anteriorly, there were slightly more TRAP+ osteo/chondroclasts present in the area where Meckel’s cartilage would normally be located (**Supplemental Fig 1C-1D**). Posteriorly, there was no difference in the number of TRAP+ cells between controls and double mutants (**Supplemental Fig 1C’-1D’**). Together these results indicate that intramembranous bone formation was not affected in double mutants, and the absence of Meckel’s cartilage was not due to accelerated resorption by increased activity of osteo/chondroclasts.

### *Prdm3* and *Prdm16* double mutants fail to form Meckel’s cartilage

With the lack of Meckel’s cartilage, anteriorly and posteriorly throughout the mandible at E15.5 and little changes in bone formation and bone resorption, we analyzed the development of the Meckel’s cartilage at earlier stages of development to see when in developmental time, this phenotype occurs. Whole mount cartilage skeletal preparations at E14.5 also showed complete absence of Meckel’s cartilage in *Prdm3^fl/fl^;Prdm16^fl/fl^;Wnt1-Cre^+/Tg^* double mutants (**Fig 2A-2B**). This was confirmed histologically with coronal sections through the mandible that were stained for Alcian blue and nuclear fast red. Meckel’s cartilage robustly stained positive for Alcian blue in control animals with the mandible primordium starting to form in the area surrounding the Meckel’s cartilage (**Fig 2A’**). In *Prdm3^fl/fl^;Prdm16^fl/fl^;Wnt1-Cre^+/Tg^* double mutant animals, Meckel’s cartilage was completely missing and the mandible primordium appears disorganized and misshapen (**Fig 2B’**). The lack of Meckel’s cartilage occurs only with loss of both *Prdm3* and *Prdm16*, as single mutants both develop Meckel’s cartilage, albeit hypoplastic in *Prdm16* single mutants (**Supplemental Fig 1A’’-C’’**). To determine if Meckel’s cartilage defects were due to changes in cell number, proliferation and apoptosis was assessed at the stage of chondroblast commitment and initiation of chondrogenic differentiation programs in the mandible primordium. We found that the absence of Meckel’s cartilage was not due to differences in neural crest cell proliferation or apoptosis after their migration into the mandibular arch, as there were no changes in the number of phosphoH3 positive cells or cleaved caspase 3 positive cells, respectfully, relative to tissue area of the mandibular prominence at E11.5 (**Fig 2C-H**).

**Figure 2.**
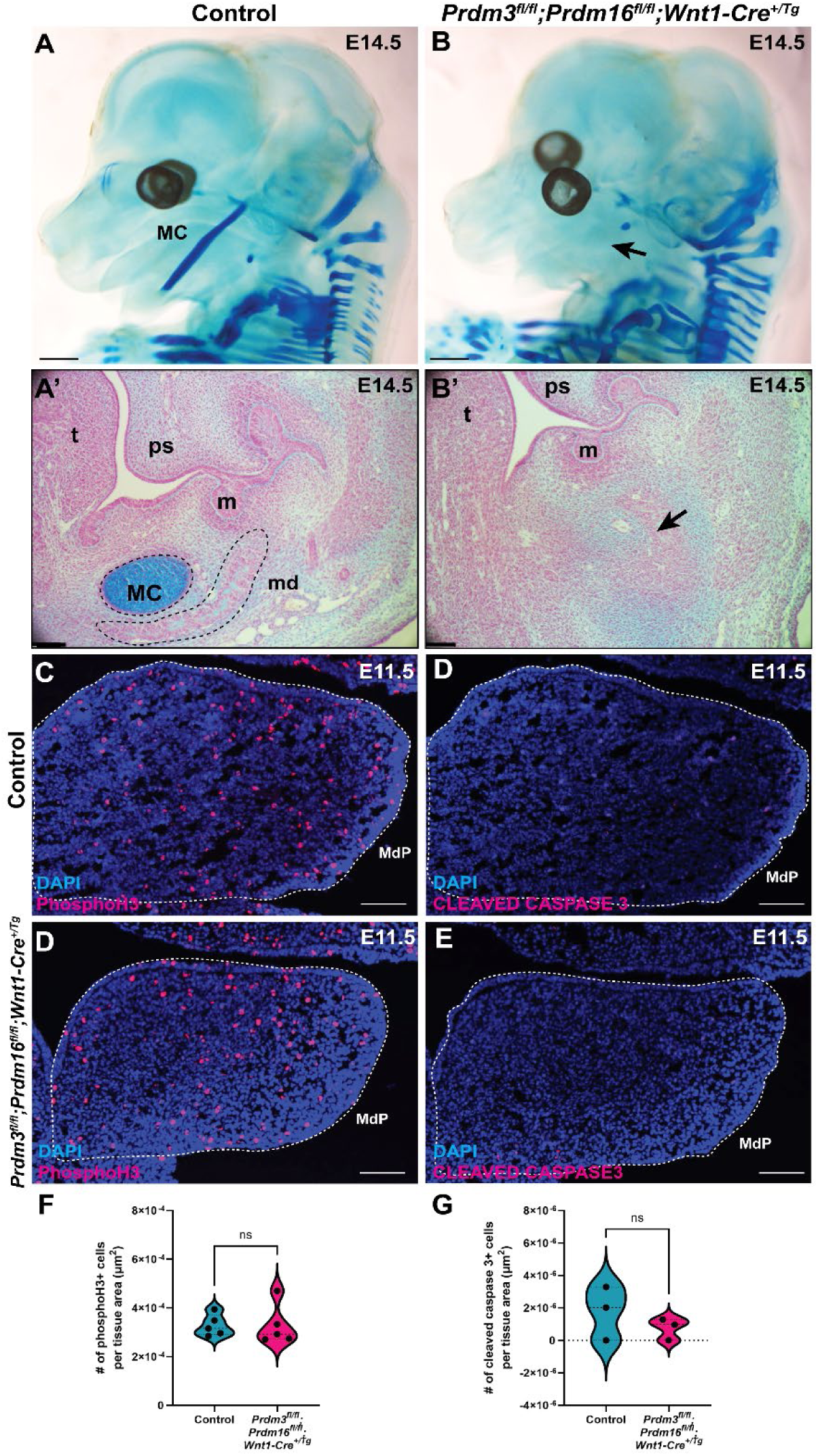
*Prdm3* and *Prdm16* are both required for the formation of Meckel’s cartilage in the developing mandibular process. (A-B) Alcian blue stained and cleared embryos at E14.5. Shown are lateral views of the whole head. Black arrow indicates absence of Meckel’s cartilage in double mutants. (A’, B’) Alcian Blue/Nuclear Red stained coronal sections through the mandible at E13.5. Black arrow again demonstrates absence of Meckel’s cartilage. (C-E) Phospho-histone H3 immunofluorescence on sections through the mandibular process at E11.5. (E) Quantification of the # of phosphoH3+ cells per tissue area (n=5 for control; n=5). (F-H) Cleaved caspase 3 immunofluorescence on sections through the mandibular process at E11.5 to detect apoptotic cells. (H) Quantification of the # of cleaved caspase 3+ cells per tissue area (n=3 for control and double mutants). Abbreviations: mc, Meckel’s cartilage; md, mandible; MdP, mandibular process; m, molar; ps, palatal shelf; t, tongue. ns, not significant (unpaired, two-tailed Student’s *t* test).

### Loss of Prdm3 and Prdm16 leads to increased Runx2+ osteoblast progenitors in the mandibular anlage

Neural crest cells populate the mandibular arch and are patterned along the dorsoventral axis through expression of a series of patterning genes including *Dlx5, Dlx6, Hand2, Ednra*, among others^26^. To assess if expression of these genes were altered in double mutants, RT-qPCR was performed on dissected mandibular processes from control or *Prdm3^fl/fl^;Prdm16^fl/fl^;Wnt1-Cre^+/Tg^*at E11.5. There was no significant difference in expression of *Dlx5*, *Dlx6*, *Hand2* or *Ednra* (**Fig 3A**), suggesting there were no neural crest patterning defects. As neural crest cells in the pharyngeal arches become specified to osteochondroprogenitor fates, they lose *Sox10* expression but continue to express *Sox9* and begin to express *Runx2*. At this point, neural crest-derived osteochondroprogenitors that are destined to become chondrocytes of the Meckel’s cartilage will repress *Runx2* to favor *Sox9* expression, which facilitates the activation of the chondrogenic program including *Col2a1*. Those osteochondroprogenitors that are instead destined to become osteoblasts will repress *Sox9* but continue to express *Runx2* and subsequently activate the osteogenic differentiation program. To assess changes to the expression of these osteochondroprogenitor differentiation genes, RT-qPCR was performed on dissected mandibular processes from control and *Prdm3^fl/fl^;Prdm16^fl/fl^;Wnt1-Cre^+/Tg^* double mutants at E11.5, the stage at which neural crest osteochondroprogenitors start their condensation events and chondrogenic commitment and differentiation begins (**Fig 3B**). While there was no drastic change in expression of chondrogenic genes (*Sox9* and *Col2a1*) in *Prdm3^fl/fl^;Prdm16^fl/fl^;Wnt1-Cre*^+/Tg^ double mutants, there was a significant increase in *Runx2* and *Osterix* (*Osx*) expression in the mandibular process of these animals (**Fig 3B**). The mandibular tissue from these animals also expresses significantly higher levels of *Sox10*. These results suggest the neural crest mesenchyme has reached a *Sox9+/Runx2*+ progenitor state, but fail to turn on the chondrogenic program and instead favors osteogenic differentiation. We did not observe any statistically significant differences one day earlier at E10.5 when the neural crest mesenchyme in the mandibular process is expanding, supporting the hypothesis that PRDM3 and PRDM16 are necessary for the osteochondroprogenitor differentiation program rather than proliferation and expansion of the neural crest pool in the mandibular arch (**Supplemental Fig 4**).

**Figure 3.**
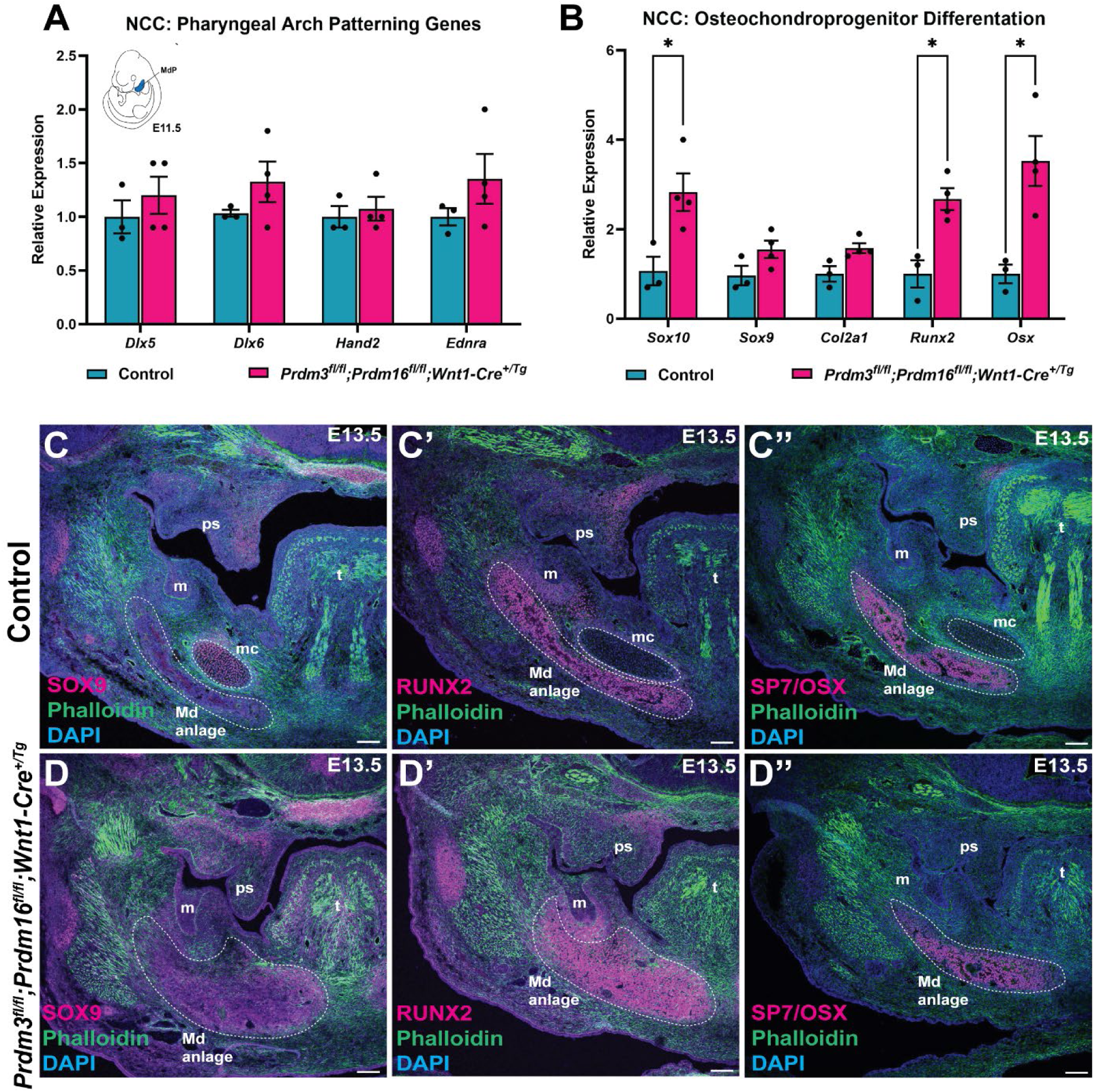
Loss of *Prdm3* and *Prdm16* causes increased Runx2+ osteochondroprogenitors in the mandible primordium. (A-B) RT-qPCR for the indicated mRNA transcripts isolated from the mandibular processes of control and *Prdm3;Prdm16* double mutants at E11.5. (C-D’’) Immunofluorescence was performed for osteochondroprogenitor program markers on coronal sections through the mandible: SOX9 (C, D), RUNX2 (C’, D’) and SP7/OSX (C’’-D’’) (shown in magenta). Sections were counterstained with phalloidin (green) and DAPI (blue). Abbreviations: mc, Meckel’s cartilage; Md anlage, mandible anlage; MdP, mandibular process; m, molar; ps, palatal shelf; t, tongue. *P≤0.05; ns, not significant (unpaired, two-tailed Student’s *t* test).

To determine if the changes in gene expression at E11.5 continued on to later time points, we validated protein expression with immunofluorescence at a later stage of development. At E13.5, SOX9 protein expression is spatially restricted to the Meckel’s cartilage chondrocytes, while RUNX2 (and subsequent SP7/OSX) expression is restricted to the mandible anlage in coronal sections of the mandible in control animals (**Fig 3C, 3C’, 3C’’**). In stage matched *Prdm3^fl/fl^;Prdm16^fl/fl^;Wnt1-Cre^+/Tg^*double mutants, Meckel’s cartilage chondrocytes were completely absent, while SOX9 protein expression was low in the mandible anlage. This low-level expression of SOX9 correlated with a strikingly large expansion of RUNX2 positive cells in this same tissue region (**Fig 3D, D3’, 3D’’**). Taken together, these results suggest that both PRDM3 and PRDM16 are necessary for NCC-derived osteochondroprogenitor differentiation programs during the formation of the developing mandible. Loss of both *Prdm3* and *Prdm16* leads to an increase in neural crest derived-RUNX2 positive osteoblast progenitors at the expense of neural crest derived SOX9 positive chondrocytes that would go on to form the Meckel’s cartilage.

### PRDM3 and PRDM16 regulate Wnt/β-catenin transcriptional activity in the mandibular process to control osteochondroprogenitor fates

To determine how PRDM3 and PRDM16 are controlling this neural crest osteochondroprogenitor fate decision in the mandible, we mined previously published H3K27me3, H3K27ac, and H3K4me1 chromatin immunoprecipitation paired with sequencing (ChIP-seq) datasets from E11.5 mouse mandibular processes (FACEBASE)^27^. Motif analysis revealed no predictive putative PRDM3 or PRDM16 binding sites at promoters of key osteochondroprogenitor cell genes (*Sox9*, *Runx2*) (**Supplemental Fig 5**). While this does not exclude the possibility that PRDM3 and PRDM16 control distal enhancer regions of these different loci, or work in large protein complexes to regulate expression of these genes, this analysis suggests that neither PRDM3 nor PRDM16 directly regulate the expression of these genes at this stage of development, but rather control these fate decisions through a secondary mechanism.

We previously showed that PRDM3 and PRDM16 spatiotemporally control Wnt/β-catenin transcriptional activity antagonistically in both zebrafish and single *Prdm3* and *Prdm16* neural crest specific mouse mutants, whereby PRDM3 serves as the repressor of gene expression while PRDM16 serves as an activator of gene expression, albeit both functioning to regulate expression of Wnt/β-catenin signaling components^20^. In long bone, SOX9 and β-catenin antagonize one another to control chondrogenesis^28^. SOX9 can compete with β-catenin for TCF/LEF binding sites to control chondrogenic genes^28^. High abundance of β-catenin is inhibitory to chondrogenesis and instead promotes osteoblastogenesis^29,30^. We hypothesized that loss of both *Prdm3* and *Prdm16* together alters expression of Wnt/β-catenin signaling components and in doing so compromises SOX9 transcriptional activity and shifts neural crest osteochondroprogenitors toward the osteoblast lineage over a chondrogenic fate (**Fig 4A**). To test this hypothesis, we performed RT-qPCR for β-catenin. mRNA transcripts of β-catenin (*Ctnnb1*) were significantly elevated in the mandibular process of *Prdm3^fl/fl^;Prdm16^fl/fl^;Wnt1-Cre^+/Tg^* mutants compared to controls at E11.5 (**Fig 4B**). To assess spatial localization, immunofluorescence for the active, non-phosphorylated form of β-catenin was performed on coronal sections through the mandible at E13.5 (**Fig 4C-D**). Protein levels of active β-catenin remained elevated in *Prdm3^fl/fl^;Prdm16^fl/fl^;Wnt1-Cre^+/Tg^* double mutants compared to control animals and localized spatially in the mandibular anlage and surrounding cells that formed in place of the Meckel’s cartilage chondrocytes (**Fig 4D**). Noticeably, the osteoblasts within this region appear more compact and less organized compared to controls. Higher magnification of osteoblasts within this region shows increased active β-catenin at the cellular junctions (**Fig 4C’-D’**). Together, these results suggest PRDM3 and PRDM16 work together to spatiotemporally control Wnt/β-catenin activity within the developing mandibular process to control neural crest derived osteochondroprogenitors fate decisions. Loss of both in the neural crest lineage leads to increased Wnt/β-catenin signaling which subsequently represses SOX9 activity, leading to an increase in RUNX2 osteoblast progenitors in the mandible anlage at the expense of Meckel’s cartilage chondrocytes.

**Figure 4.**
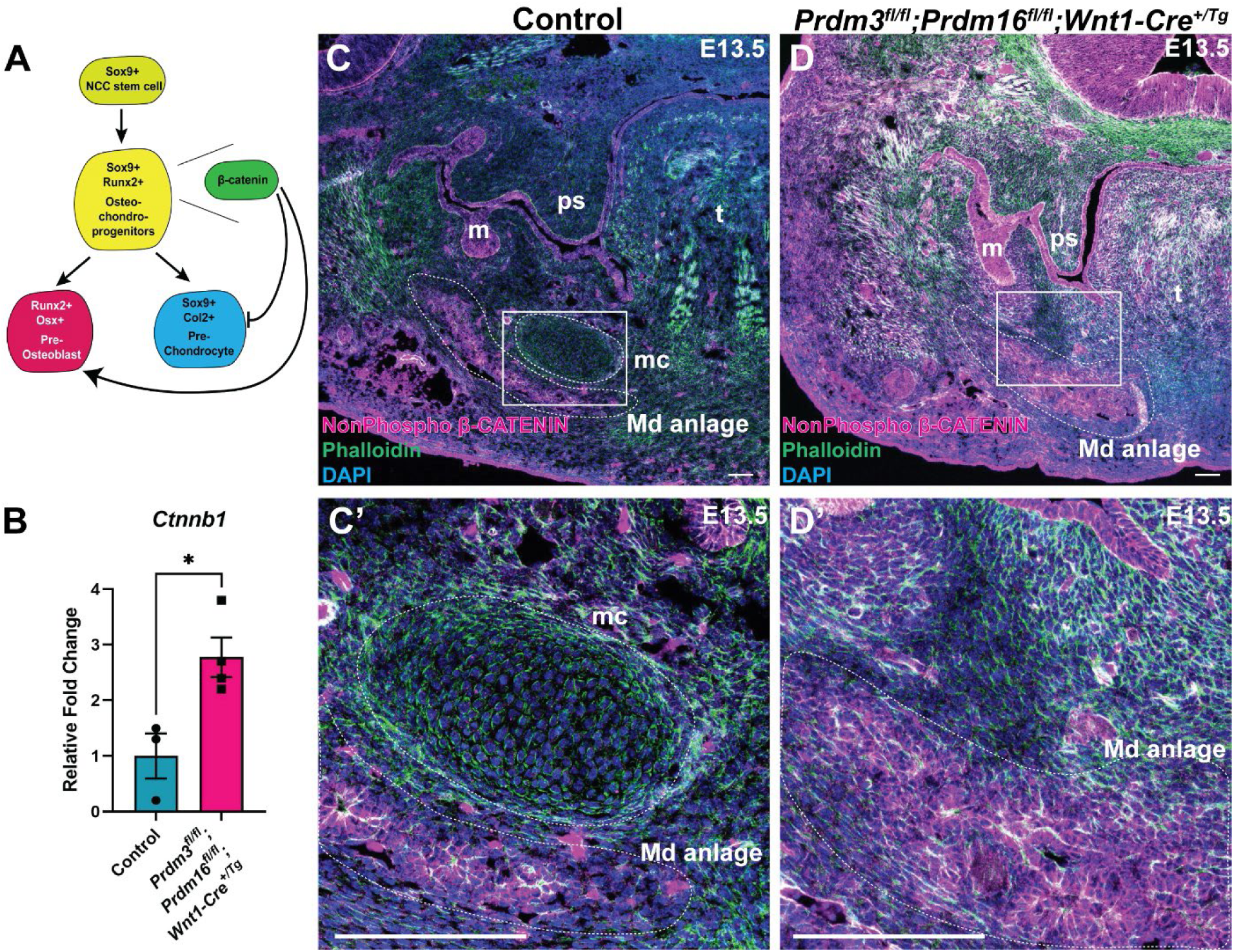
PRDM3 and PRDM16 regulate Wnt/β-catenin transcriptional activity in the mandibular process to control osteochondroprogenitor fates. (A) *Sox9*+ NCC stem cells destined to become osteochondroprogenitor cells in the craniofacial complex will co-express *Runx2*. At this point the osteoblast progenitors will continue to express *Runx2*, turn off *Sox9* and begin to express *Osx*. Chondrogenic precursors will continue to express *Sox9*, turn off *Runx2* and begin to express *Col2a1*. β-catenin directly affects the function of SOX9 and vice versa to control osteochondroprogenitor fates. (B) RT-qPCR for *Ctnnb1* on mandibular processes at E11.5. (C-D’) Immunofluorescence for non-phosphorylated (Active) β-catenin (magenta) was performed on coronal sections through the mandible at E13.5. Sections were counterstained with DAPI (blue) and phalloidin (green). (C’-D’) High magnification of the white box inlet shown in C and D. Abbreviations: mc, Meckel’s cartilage; Md anlage, mandible anlage; m, molar; ps, palatal shelf; t, tongue. *P≤0.05 (unpaired, two-tailed Student’s *t* test).

**Figure 5.**
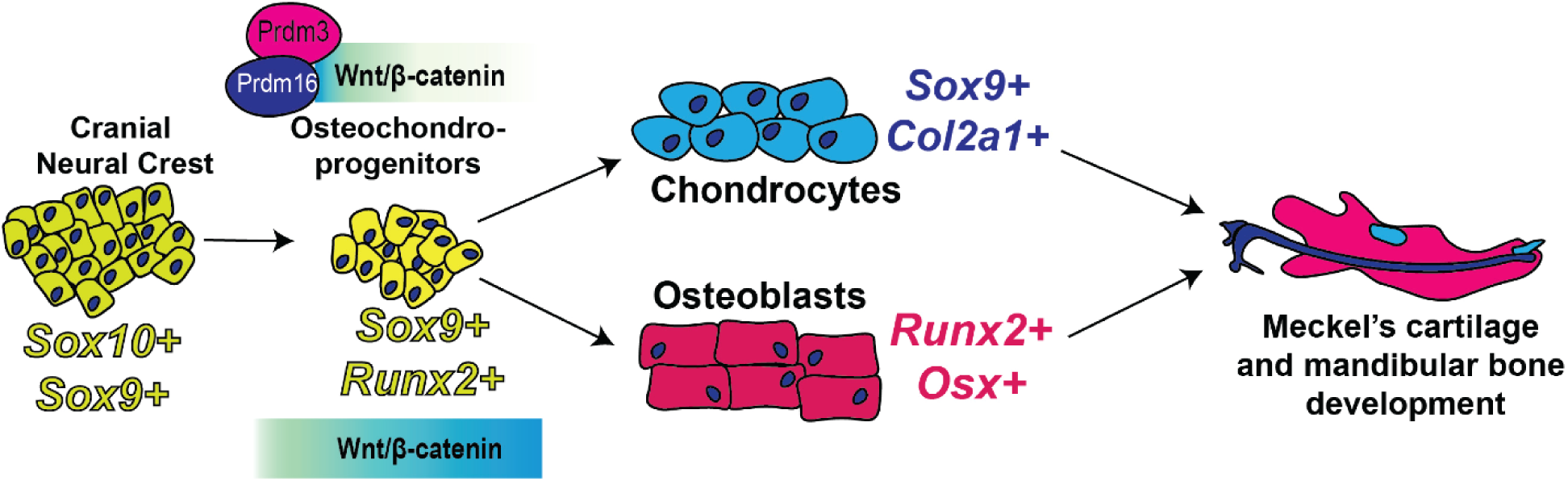
PRDM3 and PRDM16 function together to spatiotemporally restrict Wnt/β-catenin activity in cranial neural crest osteochondroprogenitors to allow for chondrogenesis during Meckel’s cartilage formation and development of the mandibular process.

## Discussion

In this study, we investigated the genetic interaction between two PRDM histone methyltransferase paralogs, *Prdm3* and *Prdm16* in mammalian cranial neural crest development and formation of craniofacial derivatives and structures, particularly in Meckel’s cartilage and mandibular bone development. We found that loss of both of these chromatin remodelers results in more dramatic craniofacial phenotypes compared to single mutants, notably a hypoplastic mandible. This mandibular hypoplasia is due in part to the complete absence of Meckel’s cartilage chondrocytes in *Prdm3;Prdm16* double mutants and reciprocally, a significant increase in RUNX2+ osteoblast progenitors. Our results suggest that PRDM3 and PRDM16 function together to restrict Wnt/β-catenin transcriptional activity to promote cranial neural crest differentiation toward chondrocytes in the mandibular process during Meckel’s cartilage formation.

We have previously shown that PRDM3 and PRDM16 are important in craniofacial cartilage by promoting chondrocyte organization during Meckel’s cartilage development, through opposing mechanisms: PRDM3 represses expression of Wnt/β-catenin signaling components, while PRDM16 activates these same transcriptional targets and that the balance of the two promote proper development of neural crest derived craniofacial structures. Given this antagonistic balance of gene expression, we were interested in assessing the genetic interaction between the two paralogs. Surprisingly, the loss of both *Prdm3* and *Prdm16* in the mammalian neural crest leads to the complete absence of Meckel’s cartilage chondrocytes in the mandibular process. Further, this is due to an overwhelming increase in Wnt/β-catenin transcriptional activity pushing these osteochondroprogenitors toward an osteoblast fate at the complete expense of the chondrogenic lineage. This suggests an epistatic hierarchal relationship between the two paralogs in relation to their spatiotemporal control of Wnt/β-catenin transcriptional activity in osteochondroprogenitor differentiation, specifically in the development of the mandibular process, and that PRDM3 may be epistatic to PRDM16 in controlling these programs at this stage of development.

In zebrafish, loss of both *prdm3* and *prdm16* leads to a rescue of craniofacial cartilage structures, hypothesized through a rebalancing of Wnt/β-catenin transcriptional activity^21^. This is not the case in the mammalian double mutants, at least in mammalian Meckel’s cartilage chondrocyte differentiation and development, where the phenotype is exacerbated with complete loss of the structure. Interestingly, this phenotype is specific only to Meckel’s cartilage, as other neural crest derived cartilages in the craniofacial complex that undergo endochondral ossification, such as the cranial base, are still present in *Prdm3;Prdm16* double mouse mutants. Perhaps this is due to the ambiguous complexity of Meckel’s cartilage, making it different from other cartilaginous structures of the craniofacial complex that would contribute to endochondral bone formation: its transient nature during development and the diverse tissue fates stemming from this structure (middle ear ossicles, dentary, and ligament)^7^. Unlike our *Prdm3;Prdm16* double mutant mice, other studies have shown the complete absence of Meckel’s cartilage with loss of key chondrogenic genes, including the conditional loss of *Sox9* in the neural crest lineage^31^ and complete knockout of *Dlx5* and *Dlx6*^32^. However, in these mouse mutants, other endochondral bones are affected including the cranial base (presphenoid and basisphenoid). This raises new questions as to why Meckel’s cartilage is specifically so phenotypically sensitive to loss of both *Prdm3* and *Prdm16*.

The mechanisms facilitating the proper gene regulatory networks and signaling events controlling the process of cranial neural crest cell fate decisions within the first pharyngeal arch remain relatively unknown. Recent single cell sequencing studies have started to elucidate the mechanisms leading to fate decisions as they form different lineages in the mandibular process and demonstrate that impaired patterning influences cranial neural crest cell fate decisions^33^. While we did not observe any striking differences in the expression of a subset mandibular patterning genes, we cannot definitively rule out the possibility that loss of the PRDM paralogs spatially disrupts patterning to then subsequently alter chondrogenic to osteogenic commitment changes. However, we do see shifts in the extrinsic signaling pathways that directly affect the activity of intrinsic factors that control cell fates, in this case the antagonism between the intrinsic and extrinsic factors controlling chondrogenesis, including the transcriptional regulator, SOX9 (intrinsic) and the signaling cascade, Wnt/β-catenin (extrinsic).

Tissue specific ablation of *Sox9* in the murine neural crest lineage leads to a complete absence of Meckel’s cartilage chondrocytes and instead cells express osteoblast markers, suggesting these cranial neural crest cells reverted to an osteoblast lineage^31^, similar to what we observed in the *Prdm3;Prdm16* double mutants. Transcriptionally active nuclear Wnt/β-catenin pathway is necessary during early mesenchymal condensation and orientation events in prechondrogenic cells, but is a potent inhibitor of chondrocyte differentiation, maturation and *Sox9* expression during skeletogenesis^29,30^.Cell culture experiments and studies in long bone axial skeletal development have suggested this antagonistic relationship between SOX9 and Wnt/β-catenin drives chondrogenesis^28^. Because Wnt ligand is detected extrinsically around chondrocyte mesenchymal condensations, yet Wnt transcriptional activity is significantly down-regulated in differentiating chondrocytes that express high levels of SOX9, SOX9 likely promotes chondrocyte differentiation and maintenance by inhibiting the anti-chondrogenic effects of Wnt/b-catenin signaling activity^28^. Further, direct interactions between SOX9 and β-catenin impact the activity of one another. For example, the N-terminal domain of SOX9 binds β-catenin and promotes its degradation while the C-terminal transactivation domain of SOX9 blocks the transcriptional activity of β-catenin, but does not lead to its degradation^34^. While a strong activator of chondrogenesis, SOX9 is also expressed in other tissues, including neural crest mesenchyme^31,34^, but whether or not this antagonistic mechanism between SOX9 and β-catenin promotes Meckel’s cartilage development in the mandibular process remains unresolved. Our results provide evidence supporting this mechanism in Meckel’s cartilage development and that from a hierarchal standpoint is regulated though the combined function of the histone methyltransferases PRDM3 and PRDM16. Specifically, we show that loss of both *Prdm3* and *Prdm16* leads to a striking increase in active β-catenin within the developing mandibular primordium which corresponds with altered SOX9 expression. Subsequently we observe an absence of chondrogenesis and instead an expansion of RUNX2+ osteoblast progenitors.

While we show that the loss of both *Prdm3* and *Prdm16* results in altered Sox9-β-catenin signaling axis, the mechanisms facilitating the requirement of both PRDM3 and PRDM16 to control this cascading event to direct the chondrogenic cell fate in Meckel’s cartilage remain unresolved. We hypothesize that each factor could be interacting in different transcriptional or repressive complexes to directly control the spatiotemporal activation or repression of these transcriptional targets. It will be interesting to further define the transcriptional and epigenetic landscape of these two paralogs during development of the mandibular arch derivatives to further support their role in maintaining osteochondroprogenitor plasticity.

In summary, we have investigated the genetic interaction between *Prdm3* and *Prdm16* in cranial neural crest development and formation of mandibular arch derivatives. We show that both of these factors are required for Meckel’s cartilage formation by controlling neural crest osteochondroprogenitor fates through spatiotemporally restricting Wnt/β-catenin transcriptional activity. Loss of these factors promotes the antagonistic relationship between β-catenin and SOX9. Ablation of both *Prdm3* and *Prdm16* leads to increased active β-catenin and a reciprocal loss of SOX9 activity which subsequently inhibits neural crest commitment towards chondrocytes, leading to the complete absence of Meckel’s cartilage. This ultimately compromises mandibular morphology and development of the Meckel’s cartilage derivatives including middle ear ossicles (tympanic ring, incus, malleus, and retroarticular process of the squamosal bone).

## Methods

### Animals

Mecom^tm1mik^ (referred to as *Prdm3^fl/fl^*)^24^, B6(SJL-Prdm16^tm.1.1Snok^/J (referred to as *Prdm16^fl/fl^*) (Jackson Laboratories), and H2afv^Tg(Wnt-Cre)11Rth^ (referred to as Wnt-Cre^+/Tg^)^23^ were all maintained on the C57/Bl6 background and housed at subthermoneutral temperatures (21-23°C) under a 12 hour light/dark cycle. Animals were provided water and food (PicoLab Rodent Diet 20) ad libitum. To generate *Prdm3fl/fl;Prdm16fl/fl* animals, *Prdm3fl/fl* mice were bred to *Prdm16fl/fl* animals to create *Prdm3fl/+;Prdm16fl/+* which were then used to generate and maintain *Prdm3fl/fl;Prdm16fl/fl* animals. For timed matings, *Prdm3fl/fl;Prdm16fl/fl* females were bred to *Prdm3fl/+;Prdm16fl/+;Wnt1-Cre^+/Tg^*males respectively. The morning a copulatory plug was observed was considered embryonic day 0.5. Embryos of matching somite numbers were used for experiments. Mice were euthanized by carbon dioxide inhalation followed by cervical dislocation as a secondary method of euthanasia. Male and female embryos were analyzed in this study and there were no sex-dependent differences in phenotypes between the two groups. Developmental stages of the embryos used are indicated in the figures and results. Mice were bred and maintained following the recommendations in the Guide for the Care and Use of Laboratory Animals of the National Institutes of Health. The protocol was approved by the University of New Mexico’s Institutional Animal Care and Use Committee.

### Genotyping

Tail clips from weanlings and tail clips or yolk sacs from embryos were lysed in DNA lysis buffer (10 mM Tris-HCl (pH 8.0), 100 mM NaCl, 10 mM EDTA (pH 8.0), 0.5% SDS) and 100 ug of Proteinase K overnight at 55°C. Genomic DNA was isolated following phenol/chloroform extraction. DNA pellets were air dried and resuspended in nuclease-free water. Genotyping for *Prdm3*, *Prdm16* and *Wnt1-Cre* alleles was performed as previously described^13,20^.

### Whole Mount Skeletal Preparations

Alcian blue and Alizarin red staining was performed as previously described for E18.5 embryos^13,35^. Briefly, mouse embryos were harvested at E18.5 in 1XPBS. Skin was removed and internal organs were eviscerated. Specimens were then fixed overnight in 95% ethanol at room temperature followed by an incubation in 100% acetone for 2 days at room temperature. Embryos were stained in Alcian blue/Alizarin red staining solution (0.015% Alcian blue, 0.05% Alizarin red, 5% Glacial acetic acid and 70% ethanol) for 3 days at 37°C. Stained embryos were rinsed in water before undergoing an initial clearing in 1% KOH overnight at room temperature. Specimens were then passed through a gradient series of decreasing KOH concentrations and increasing glycerol concentrations. Skeletal preparations were stored in 80% glycerol and imaged on either a Leica stereomicroscope with the LASX v4.4 software or with a NIKON SLR 60mm camera.

For quantification of anterior mandibular hypoplasia, Alcian Blue and Alizarin Red-stained mandibles were removed and imaged. Using ImageJ software, the anterior, posterior and total length of the mandible was measured. The ratio of the anterior portion relative to the posterior portion was quantified.

### Histology

For Alcian Blue/Eosin staining, mouse embryos were collected at the indicated time points in 1xPBS. Whole heads were dissected and removed from the animals and fixed in 4% paraformaldehyde, dehydrated through a graded ethanol series, embedded in paraffin and sectioned to a thickness of 8 μm onto glass slides. After deparaffinization and rehydration, sections were stained with 0.3% Alcian Blue for 30 sections followed by 0.05% Fast Green for 15 minutes followed by a quick rinse in1% acetic acid, then 0.1% Safranin O. Sections were dehydrated and mounted with Permount (Electron Microscopy Sciences).

For Von Kossa staining, mouse embryos were collected at the indicated time points in 1xPBS. Whole heads were dissected and removed and fixed in 4% paraformaldehyde, dehydrated through a graded ethanol series, embedded in paraffin and sectioned to a thickness of 8 µm onto glass slides. After deparaffinization and rehydration to water, sections were covered with 1% Silver Nitrate and exposed to bright light for 30 minutes. Sections were washed briefly in water then covered in 2.5% Sodium Thiosulphate and incubated at room temperature fo 5 minutes. Following another brief wash in water, sections were counterstained with Nuclear Fast Red for 5 minutes at room temperature, quickly rinsed in water and then immediately dehydrated and cleared. Sections were then mounted with Permount (Electron Microscopy Sciences) and imaged on a Leica Stereomicroscope with the LASX v4.4 software.

For Tartrate Resistant Acid Phosphatase (TRAP) staining, mouse embryos were collected at the indicated time points in 1xPBS. Whole heads were dissected and removed and fixed in 4% paraformaldehyde, dehydrated through a graded ethanol series, embedded in paraffin and coronal sections were cut to a thickness of 8 µm and collected onto glass slides. After deparaffinization and rehydration to water, sections were incubated in 10 mM Tris-HCl buffer (pH 9.0) at 37°C for 45 minutes. Sections were quickly rinsed in water then incubated in TRAP stain (Pararosaniline, Naphthol AS-TR phosphate) for 40 minutes at 37°C. Samples were rinsed with water then counterstained with 0.05% Fast Green, dehydrated and mounted with Permount.

### Immunofluorescence

For immunofluorescence, mouse embryos were collected at the indicated time points in 1xPBS. Whole heads were dissected and removed and fixed in 4% paraformaldehyde, cryoprotected in 30% sucrose/1xPBS and embedded in OCT embedding medium (TissueTek). Coronal sections at 8 µm thickness were collected and mounted on glass slides with a Leica CM1520 cryostat. Sections were rehydrated for 30 minutes in 1xPBS and then subjected to antigen retrieval using 1% SDS in 1xPBS for 5 minutes at room temperature. Sections were washed 3×5 minutes in 1xPBS then incubated in blocking solution containing 2% goat serum, 1% Bovine Serum Albumin, 1% DMSO, 0.1% triton-100 for 1 hour at room temperature. Sections were then incubated in primary antibodies diluted at 1:100 [Sox9 (Cell Signaling Technology Cat#82630, RRID:AB_2665492); Runx2, (Cell Signaling Technology Cat#12556, RRID:AB_2732805); OSX/Sp7 (Santa Cruz, Cat#sc-393060); Non-Phosphorylated (Active) β-catenin (Ser33/37/Thr41), Cat#4270); phospho-Histone H3 (Cell Signaling Technology Cat#9701, RRID:AB_331535), Cleaved Caspase 3 (Cell Signaling Technology Cat#9661, RRID:AB_2341188)] overnight at 4C. The next day, sections were washed in 1xPBS with 0.0625% TritonX-100 3×10minutes at room temperature followed by an incubation in corresponding secondary antibodies (either Goat anti-Rabbit 594, or Goat anti-Mouse 594 and phalloidin-488) diluted in blocking buffer at 1:500 for 2 hours at room temperature. Following this, sections were washed in the dark at room temperature in 1xPBS with 0.0625% TritonX-100 3×10 minutes. Sections were then incubated in DAPI (1:10000 dilution in 1xPBS) for 15 minutes at room temperature, subsequently washed in 1xPBS for 10 minutes at room temperature, mounted in Vectashield mounting medium, sealed and imaged on a Leica SP8 confocal or Zeiss 980 confocal with Airyscan.

### MicroCT analysis

MicroCT analysis was performed using a SCANCO µCT50 at the University of Colorado Anschutz Medical Campus. The microCT images were acquired from E18.5 embryos with the x-ray source set to 70kVp and 200µA. The data were collected at a resolution of 15.2 µm. 3D reconstructions were generated using Dragonfly 3D World Version 2024.1 (Comet Technologies Canada Inc.). The mandible was manually segmented using Dragonfly. MicroCT scans were uploaded to Dragonfly as Dicom files. Background noise and other bones not included in the study such as neck vertebrae were manually removed using editor tools in Dragonfly (threshold, contrast, cropping). The mandible was isolated and labeled using Dragonfly’s semiautomatic segmentation editor tool and manually using threshold levels that allowed only the mandible to be labeled. Volumetric data and Surface Area data were determined using Dragonfly’s algorithm tools. Mean measurements were compared across controls and double mutants.

### RT-qPCR

Mandibular processes were dissected on ice from 3-4 independent replicates of E11.5 *Prdm3^fl/fl^;Prdm16^fl/fl^;Wnt1-Cre^+/Tg^* double mutants or controls (*Prdm3^fl/fl^;Prdm16^fl/fl^*) embryos. Mandibular processes were lysed in TRIzol LS (Invitrogen/Life Technologies) and total RNA was isolated from these samples using a chloroform extraction. For RT-qPCR, total RNA was isolated as described and 0.5-1.0 ug of RNA was reverse transcribed to cDNA with SuperScript III First-Strand Synthesis cDNA kit (Invitrogen/Life Technologies) for real-time semi quantitative PCR (RT-qPCR) with primers (**Table S1**) and SYBR Green MasterMix (BioRad). Transcript levels were normalized to the reference gene, *Actb*. Transcript abundance and relative gene expression were quantified using the 2^-ΔΔCt^ method relative to control.

### Statistical Analysis

Data shown are the means ± SEM from the number samples or experiments indicated in the figure legends. All assays were repeated at least three times with independent samples. *P* values were determined with Student’s *t* tests.

## Acknowledgements

We thank members of the Artinger and Shull laboratories for their thoughtful discussions on this project. We thank Dr. Karin Payne and members of the CU Orthopedics department for their assistance with the MicroCT scanner. We would also like to thank Drs. Susumu Goyama and Mineo Kurokawa for the *Prdm3* mouse line. We thank the mouse facility staff at UNM for excellent animal care. Funding was provided by the National Institute of Dental and Craniofacial Research (R01 DE024034 to K.B.A. and K99/R00 DE031349 to L.C.S.).

## Author Contributions

Conceptualization: K.B.A., L.C.S.; Methodology: K.B.A., L.C.S.; Formal Analysis: Q.D.C, L.C.S.; Investigation: Q.D.C, B.V.S., L.C.S; Writing – original draft: L.C.S.; Writing – review & editing: Q.D.C., B.V.S., K.B.A., L.C.S.; Supervision: K.B.A., L.C.S.; Funding acquisition: K.B.A, L.C.S.

## References

1. Bronner, M. E. & LeDouarin, N. M. Development and evolution of the neural crest: An overview. Dev. Biol. 366, 2–9 (2012).

2. Simões-Costa, M. & Bronner, M. E. Establishing neural crest identity: a gene regulatory recipe. Development 142, 242–257 (2015).

3. Betancur, P., Bronner-Fraser, M. & Sauka-Spengler, T. Assembling neural crest regulatory circuits into a gene regulatory network. Annu. Rev. Cell Dev. Biol. 26, 581–603 (2010).

4. Cordero, D. R. et al. Cranial neural crest cells on the move: their roles in craniofacial development. Am J Med Genet A 155A, 270–279 (2011).

5. Minoux, M. & Rijli, F. M. Molecular mechanisms of cranial neural crest cell migration and patterning in craniofacial development. Development 137, 2605–2621 (2010).

6. Green, S. A., Simoes-Costa, M. & Bronner, M. E. Evolution of vertebrates as viewed from the crest. Nature 520, 474–482 (2015).

7. Svandova, E., Anthwal, N., Tucker, A. S. & Matalova, E. Diverse Fate of an Enigmatic Structure: 200 Years of Meckel’s Cartilage. Front Cell Dev Biol 8, 821 (2020).

8. Anthwal, N., Joshi, L. & Tucker, A. S. Evolution of the mammalian middle ear and jaw: adaptations and novel structures. J Anat 222, 147–160 (2013).

9. Shull, L. C., Artinger, K. B. & Lomeli Shull, C. C. Epigenetic regulation of craniofacial development and disease. (2023) doi:10.1002/bdr2.2271.

10. Di Zazzo, E., De Rosa, C., Abbondanza, C. & Moncharmont, B. PRDM Proteins: Molecular Mechanisms in Signal Transduction and Transcriptional Regulation. Biol. Basel 2, 107–141 (2013).

11. Fog, C. K., Galli, G. G. & Lund, A. H. PRDM proteins: important players in differentiation and disease. Bioessays 34, 50–60 (2012).

12. Hohenauer, T. & Moore, A. W. The Prdm family: expanding roles in stem cells and development. Development 139, 2267–2282 (2012).

13. Shull, L. C. et al. The conserved and divergent roles of Prdm3 and Prdm16 in zebrafish and mouse craniofacial development. Dev Biol 461, 132–144 (2020).

14. Warner, D. R., Wells, J. P., Greene, R. M. & Pisano, M. M. Gene expression changes in the secondary palate and mandible of Prdm16(-/-) mice. Cell Tissue Res 351, 445–452 (2013).

15. Warner, D. R. et al. PRDM16/MEL1: A novel Smad binding protein, expressed in murine embryonic orofacial tissue. Biochim. Biophys. Acta-Mol. Cell Res. 1773, 814–820 (2007).

16. Bjork, B. C., Turbe-Doan, A., Prysak, M., Herron, B. J. & Beier, D. R. Prdm16 is required for normal palatogenesis in mice. Hum Mol Genet 19, 774–789 (2010).

17. Jugessur, A. et al. Maternal genes and facial clefts in offspring: a comprehensive search for genetic associations in two population-based cleft studies from Scandinavia. PLoS One 5, e11493 (2010).

18. Shaffer, J. R. et al. Genome-Wide Association Study Reveals Multiple Loci Influencing Normal Human Facial Morphology. PLoS Genet 12, e1006149 (2016).

19. White, J. D. et al. Insights into the genetic architecture of the human face. Nat Genet 53, 45–53 (2021).

20. Shull, L. C. et al. PRDM paralogs antagonistically balance Wnt/beta-catenin activity during craniofacial chondrocyte differentiation. Development 149, (2022).

21. Baizabal, J. M. et al. The Epigenetic State of PRDM16-Regulated Enhancers in Radial Glia Controls Cortical Neuron Position. Neuron 99, 239–241 (2018).

22. He, H. et al. PRDM3/16 regulate chromatin accessibility required for NKX2-1 mediated alveolar epithelial differentiation and function. Nat. Commun. 15, 8112 (2024).

23. Danielian, P. S., Muccino, D., Rowitch, D. H., Michael, S. K. & McMahon, A. P. Modification of gene activity in mouse embryos in utero by a tamoxifen-inducible form of Cre recombinase. Curr Biol 8, 1323–1326 (1998).

24. Goyama, S. et al. Evi-1 is a critical regulator for hematopoietic stem cells and transformed leukemic cells. Cell Stem Cell 3, 207–220 (2008).

25. Hoyt, P. R. et al. The Evi1 proto-oncogene is required at midgestation for neural, heart, and paraxial mesenchyme development. Mech Dev 65, 55–70 (1997).

26. Santagati, F. & Rijli, F. M. Cranial neural crest and the building of the vertebrate head. Nat. Rev. Neurosci. 4, 806–818 (2003).

27. Visel, A. et al. ChIP-seq of multiple histone marks and RNA-seq from e11.5 mouse face subregions. FaceBase Consort. (2017).

28. Topol, L., Chen, W., Song, H., Day, T. F. & Yang, Y. Sox9 Inhibits Wnt Signaling by Promoting-Catenin Phosphorylation in the Nucleus * □ S. (2009) doi:10.1074/jbc.M808048200.

29. Hwang, S. G. et al. Regulation of beta-catenin signaling and maintenance of chondrocyte differentiation by ubiquitin-independent proteasomal degradation of alpha-catenin. J Biol Chem 280, 12758–12765 (2005).

30. Ryu, J. H. et al. Regulation of the chondrocyte phenotype by beta-catenin. Development 129, 5541–5550 (2002).

31. Mori-Akiyama, Y., Akiyama, H., Rowitch, D. H. & de Crombrugghe, B. Sox9 is required for determination of the chondrogenic cell lineage in the cranial neural crest. Proc Natl Acad Sci U A 100, 9360–9365 (2003).

32. Robledo, R. F., Rajan, L., Li, X. & Lufkin, T. The *Dlx5* and *Dlx6* homeobox genes are essential for craniofacial, axial, and appendicular skeletal development. Genes Dev. 16, 1089– 1101 (2002).

33. Yuan, Y., et al. Spatiotemporal Cellular Movement and Fate Decisions during First Pharyngeal Arch Morphogenesis. Sci. Adv vol. 6 119–135 https://www.science.org (2020).

34. Akiyama, H. et al. Interactions between Sox9 and -catenin control chondrocyte differentiation.

35. Wallin, J. et al. The role of Pax-1 in axial skeleton development. Development 120, 1109–1121 (1994).

